# Occurrence and persistence of pseudo-tail spots in the barn swallow

**DOI:** 10.1101/2024.03.06.583633

**Authors:** Masaru Hasegawa

## Abstract

While numerous studies have confirmed sexual selection for ornamental traits in animals, it remains unclear about how animals exaggerate ornamentation beyond their original traits. I found that some Asian barn swallows *Hirundo rustica gutturalis* possessed pseudo-tail spots in their undertail coverts. A close inspection showed its remarkable resemblance to the white tail spots, a well-known sexual signal in this species, although pseudo-tail spots in the undertail covers do not incur any flight cost, unlike the white tail spots on the tail itself. Presence of pseudo-tail spots can thus represent an initial stage of a deceptive elaboration as predicted by sexual selection theory. The frequency of pseudo-tail spots in the study population remained low even a decade after the first observation (ca. 7%), but was higher compared to other populations (e.g., 1% in another Japanese population). The slow progress of evolution, perhaps due to the low detectability of the trait, provides a unique opportunity to observe contemporary evolution of ornament exaggeration across traits.

---

Animals often exhibit ornamental traits that appear to have little viability function. Since Darwin first proposed the idea of sexual selection (Darwin 1859, 1871), numerous empirical studies have revealed that animal ornamentation is, in fact, sexually selected through mate preference and intrasexual contest (reviewed in Andersson 1994; Hill 2006; Brooks and Griffith 2010). One prevailing explanation for sexual selection posits that ornaments evolve because they convey information of signalers and thus signal receivers benefit from paying attention to them (i.e., honest signal; reviewed in Griffith and Pryke 2006). Although empirical studies elucidate the evolution of specific ornamental traits (e.g., tail length; Møller 1994; see also Svensson and Gosden 2007 for contemporary evolution of these ornaments), many animals exhibit gaudy, complex ornamentation ACROSS multiple traits (e.g., peacock’s long train adorned with multicolored eyespots: Dakin and Montgomerie 2013). It remains elusive about how animals exaggerate ornamentation beyond the original, rudimentary form into extreme, elaborated expressions (e.g., via honest signaling or other processes).

Theoretically, ornaments can be elaborated through sexual conflict. Hill (1994) theorized that once female mate preference for a costly signal spreads in a population, there will be strong selection on males to express the display character, sometimes novel traits, at reduced cost regardless of their actual quality (see also van Doorn and Weissing 2006). This “deceptive” trait of signalers (sensu Hill 1994; see also Bradbury and Vehrencamp 1998) can rapidly spread through a population, which allows all individuals to converge on the maximum expression. However, the rapid fixation makes it challenging to observe the elaboration process within a single species, necessitating reliance on phylogenetic comparative analysis (Hill 1994).

Here, I found the potential evolution of such deceptive elaboration in the Asian barn swallow, *Hirundo rustica gutturalis*. In barn swallows, the size of white tail spots is related to several indices of mate preference, including early breeding onset, high within-pair paternity, and differential maternal investment, indicating intersexual selection for the ornament (Kose and Møller 1999; Kose et al. 1999; Hasegawa et al. 2010, 2012; reviewed in Hasegawa 2018). A recent meta-analysis shows that white tail spots are especially important in the Asian subspecies compared to the European subspecies *H. r. rustica* (Romano et al. 2017). This is further corroborated by the finding that the Asian subspecies exhibit greater sexual dimorphism in the size of white tail spots compared to European subspecies (Hasegawa et al. 2017). At the same time, white tail spots exhibit inherent vulnerability to feather breakage because of structural weakness as they lack melanin pigments that strengthen feathers (Kose and Møller 1999) and because of feather damage caused by feather lice, *Hirundoecus malleus*, which preferentially feeds on white feathers (Kose et al. 1999). Breakage of flight feathers (i.e., tail feathers) is more prevalent in individuals of lower quality (i.e., shorter-tailed males with smaller white tail spots) and this damage reduces aerodynamic efficiency (Kose and Møller 1999; Kose et al. 1999; Barbosa et al. 2002), indicating that large white tail spots are an honest signal of male quality, partially mediated by its influence on flight performance (Kose and Møller 1999; Kose et al. 1999). Saino et al. (2015) demonstrated the condition-dependence of the size of white tail spots, further supporting its role as a reliable quality indicator.

I report here that some barn swallows exhibit pseudo-tail spots in the undertail coverts and that the presence of pseudo-tails spots has been persisted in the population for a decade since the first discovery. Notably, these spots are not located on flight feathers and, therefore, do not incur any flight cost. Potential adaptive and nonadaptive explanations for the evolution and maintenance of this novel ornament and its implication in the elaboration process of ornamentation are discussed.

## METHODS

During January 19–20, 2014, I captured and ringed a total of 28 adult barn swallows (14 males and 14 females) at night, using sweep nets at their roost in Miyazaki City, Miyazaki Prefecture, Japan (31°54′N, 131°25′E). The sex of swallows was determined from morphological measurements (e.g., males have longer tails; see Turner 2006). Among the 28 captured adults, two individuals exhibited pseudo-tail spots on their undertail coverts, which are typically whitish in Asian barn swallows (see Hasegawa 2005 for detailed feather coloration in a breeding population at Niigata Prefecture). Unfortunately, I paid little attention in the first individual at the capture (resultantly, I failed to make a note to identify the individual and take photographs), and thus I focused on describing the second individual in greater detail. To investigate the change in the frequency of pseudo-tail spots ten years later, I conducted a follow-up survey in the same population during February 2023 and 2024 (specifically, February 12-17, 2023, and February 14-18, 2024), employing the same methodology used in the initial observation in 2014 (see above). This follow-up survey aimed to test the prediction of rapid fixation of deceptive ornamentation, expecting an increase in the frequency of pseudo-tail spots (see Introduction). During 2024, two females out of 17 captured adult swallows exhibited pseudo-tail spots (but none of the 15 adult swallows captured in 2023 exhibited pseudo-tail spots).

Since the survey in Miyazaki Prefecture involved a wintering population (possibly including some migrants), I conducted an additional field survey during the breeding season (March–May) of 2014–2015 in a breeding population around Hayama-cho, Kanagawa Prefecture, Japan (35°16′N, 139°35′E; see Hasegawa et al. 2017 for a detailed explanation of the study population). The sex of swallows was determined from morphological measurements. During 2014 and 2015, one female and one male, among the 110 and 103 captured adult swallows, respectively, exhibited pseudo-tail spots on their undertail coverts.

## RESULTS

Unlike typical males that have whitish undertail coverts, which obscure the white spots on the tail (Fig. 1, upper panel), the focal male exhibited undertail coverts that closely resembled the white tail spots on the tail (i.e., “pseudo-tail spots”; Fig. 1, middle panel). In barn swallows, the two central tail feathers have no white spots (Fig. 1, lower panel), which may be effective for flashing the white spots on their other tail feathers when spread (sensu JabŁoński 1999). Pseudo-tail spots can conceal the black central black tail feathers, increasing the number (and thus total area) of white spots, when spread tails are viewed from below (see the difference between Fig. 1, middle and lower panels). The proportion of individuals with pseudo-tail spots in the population was 7.1% (Table 1; see Methods). Close-up pictures of the undertail covert feathers of the focal individual alongside ten additional swallows (five males and five females) captured during the same season in the same population can be found in Fig. S1, showing a notable difference between feathers with and without pseudo-tail spots.

**Table 1.** Frequency of individuals with and without pseudo-tail spots in the Miyazaki population of the Asian barn swallow.

|  | 2014 | 2023 + 2024 | Total |
| --- | --- | --- | --- |
| With pseudo-tail spots | 2 (1?) | 2 (2) | 4 (3?) |
| Without pseudo-tail spots | 26 (13?) | 28 (14) | 58 (27?) |
| Total | 28 (14) | 30 (16) | 62 (30) |

Data from 2023 and 2024 were combined due to small sample sizes (2023: 0 out of 15 (7), 2024: 2 (2) out of 17 (9); note that two males were recaptured across years and thus total number of different individuals were 30). Figures in parentheses indicate females (note that one individual with pseudo-tail spots in 2014 was failed to be identified in 2014; see Methods).

**Fig. 1.**
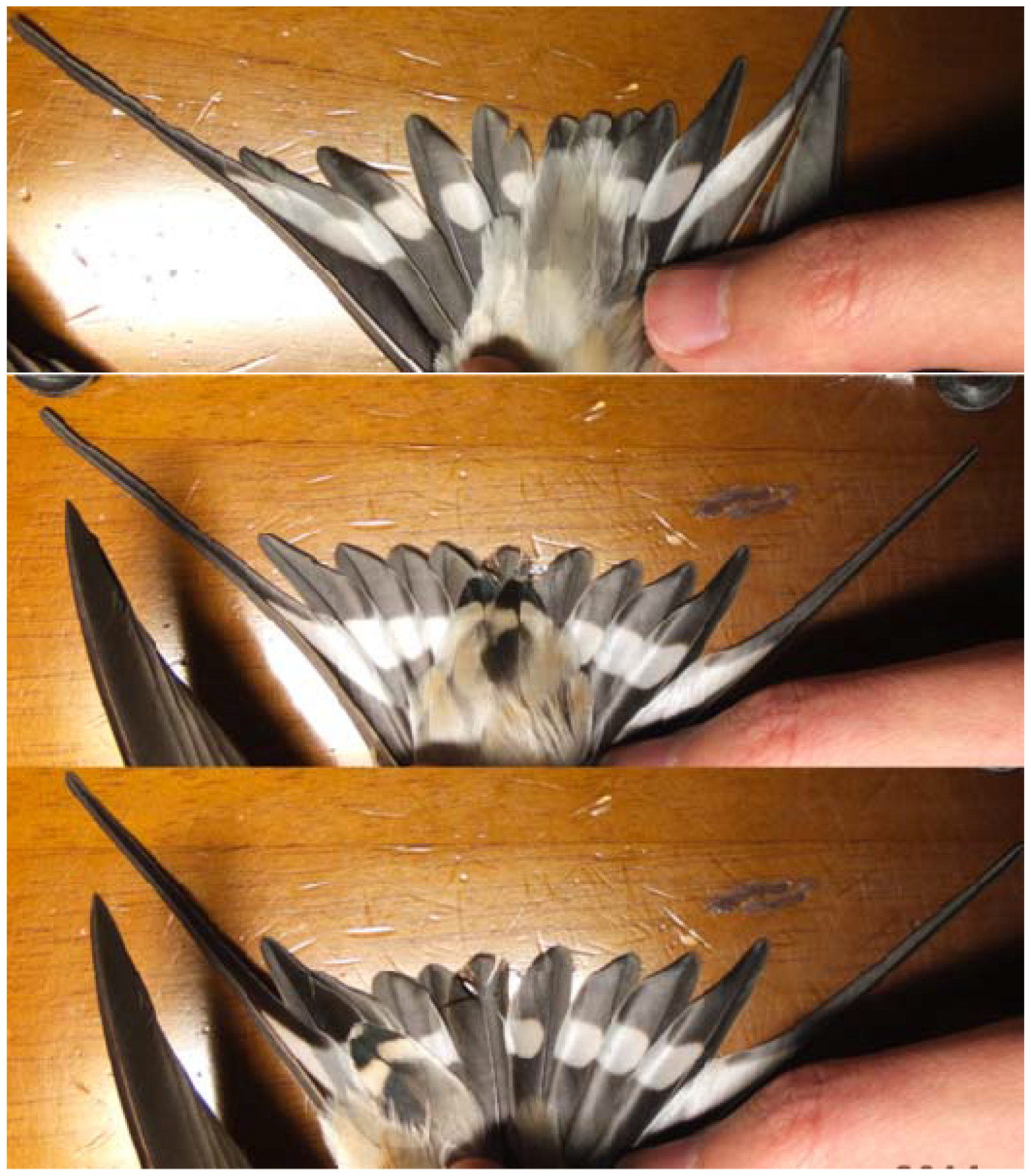
Pseudo-tail spots found in a male Asian barn swallow, *Hirundo rustica gutturalis* in the Miyazaki prefecture in 2014 (upper panel: undertail coverts of typical male; middle panel: pseudo-tail spots found in undertail coverts of the focal male; lower panel: pseudo-tail spots were moved aside, revealing the central tail feathers that lack white spots).

Ten years (2023–2024) after the initial 2014 observation of individuals with pseudo-tail spots, I conducted a follow-up survey in the same population. I found two females exhibiting pseudo-tail spots in this period (Table 1; Fig. 2; also see Fig. S2 right panel for another female). The proportion of individuals with pseudo-tail spots in this period was 6.7%, which was not significantly different from that observed in 2014 (Fisher’s exact test, P = 1.00).

**Fig. 2.**
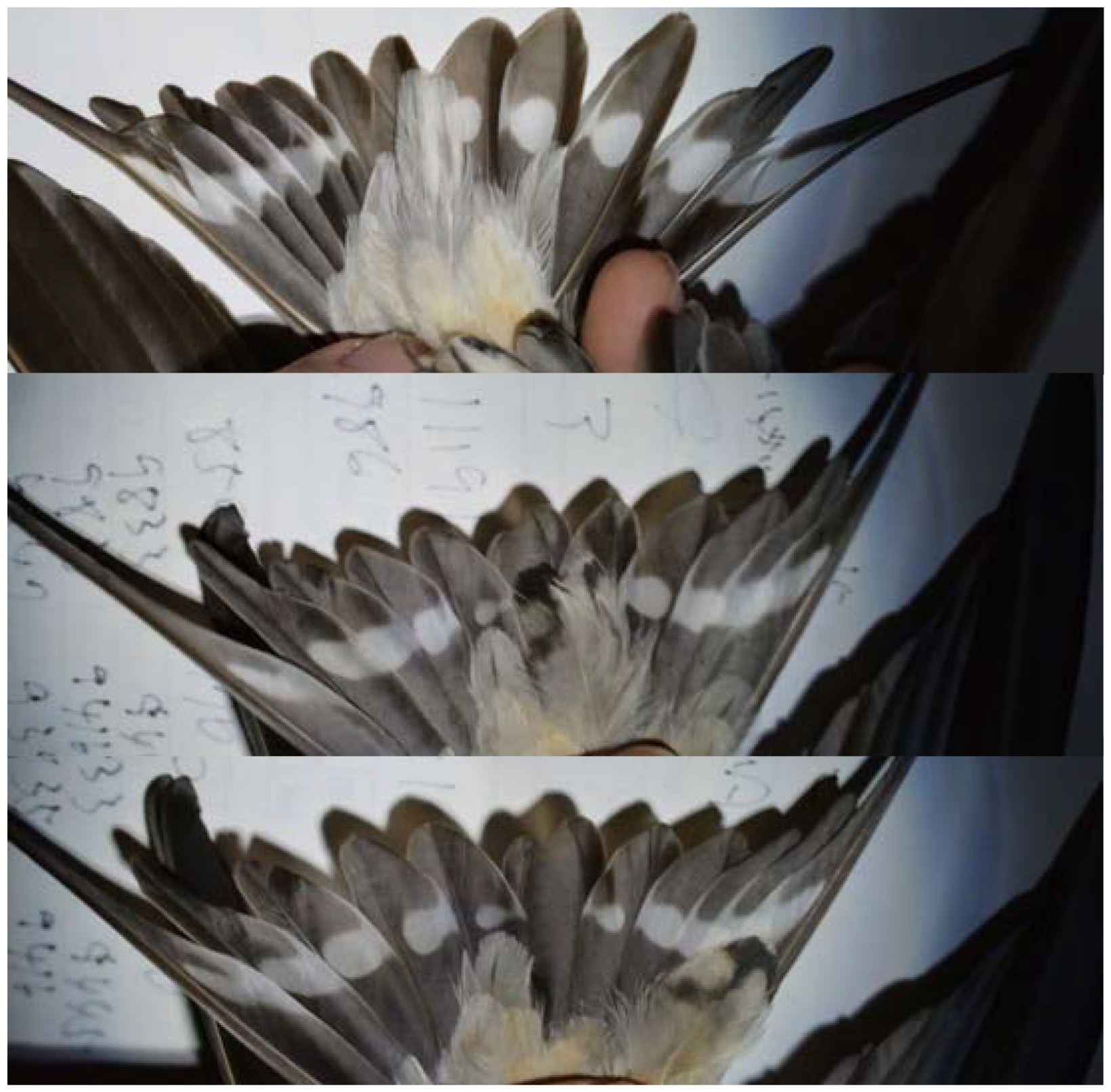
Pseudo-tail spots found in a female Asian barn swallow, *Hirundo rustica gutturalis* in the Miyazaki Prefecture in 2024 (upper panel: undertail coverts of typical female; middle panel: pseudo-tail spots found in undertail coverts of the focal male; lower panel: pseudo-tail spots were moved aside, revealing the central tail feathers that lack white spots).

Pseudo-tail spots were found in the breeding population of Hayama-cho in Kanagawa prefecture, too. During a capture survey in 2014, one female (out of 110 individuals; 0.9%) exhibited pseudo-tail spots (Fig. S3). In 2015, one male (out of 103 individuals; 1.0%) exhibited the same feature (Fig. S2 left panel). The frequency of individuals with pseudo-tail spots in the breeding population did not differ significantly between 2014 and 2015 (Fisher’s exact test, P = 1.00).

After pooling all individuals in each of the two populations, frequency of pseudo-tail spots was significantly higher in the Miyazaki population (6.9%; Table 1) compared to the Hayama population (1.0%; male: 1/96; females: 1/94; Fisher’s exact test, P = 0.038; note that we counted the number of different individuals to avoid pseudoreplication).

## DISCUSSION

The main finding of the current study is the occurrence and persistence of pseudo-tail spots in undertail coverts that closely resemble white tail spots, in the Miyazaki population of Asian barn swallows. Although black shaft streaks, tips, or spots on undertail coverts have been reported in barn swallows (e.g., Pedler 1977; Brombach 1984; reviewed in Cramp 1988; note that these are not white tail spots-like shape), the current study presents the first record of pseudo-tail spots, in which white spots distinctly appear against the black background of the undertail coverts. Possible explanations for the occurrence of pseudo-tail spots include (1) genetic misexpression, i.e., genes responsible for tail feather pigmentation pattern could become mistakenly expressed in the undertail coverts; (2) gene introgression from other species (e.g., interbreeding with other hirundine species, such the Pacific swallows *Hirundo tahitica*, which have black undertail coverts, might introduce these genes); and (3) random mutation, i.e., de novo mutations causing melanization in typically white undertail coverts. Although I could not distinguish these and possible other causations, environmental factor alone (e.g., stains) cannot adequately explain the complex (and consistent) feather patterning, suggesting the genetic control of this novel trait, which is further reinforced by its persistence in the population a decade after its initial observation. Because white tail spots in Asian barn swallows serve as sexual signals (reviewed in Romano et al. 2017), the emergence of pseudo-tail spots can be regarded as the initial stage of the elaboration process of ornamentation beyond the original trait (i.e., the “t3” stage in fig. 1 in Hill 1994; note that this does not necessarily mean immediate viability or sexual functions of the novel trait; see below).

The occurrence and persistence of a new trait do not always imply it is adaptive. In addition to several adaptive explanations including sexual selection (see Introduction), nonadaptive mechanisms, such as genetic drift and hitchhiking, could explain the persistence of neutral or even slightly maladaptive traits (Bergrstom & Dugatkin 2016) and it remains unclear about whether barn swallows inherit pseudo-tail spots across generations from the current study (i.e., its heritability remains untested). However, the population persistence for a decade, together with population difference in the frequency (see results), indicates that they would not be under a complete random process. This is further reinforced by the absence of similar observations in the nominate subspecies, despite extensive research on this well-studied model system in the field of behavioral ecology (reviewed in Møller 1994; Turner 2006). It is premature to conclude whether or not the observed pattern is maintained by selection, but it is likely that genetic mechanism that caused pseudo-tail spots (or, in other words, failure to maintain typically whitish undertail coverts) could potentially become a target of selection.

Given that the fixation of additional ornamental traits is expected to be a rapid process (Hill 1994), it is intriguing to see the low frequency of pseudo-tail spots even after ten years. This is particularly curious, considering that sexual selection on white tail spots are intense in the Asian subspecies (Romano et al. 2017), and that pseudo-tail spots incur negligible flight cost, unlike tail feathers (Gill 2007; see Introduction). However, undertail coverts cannot be seen from above due to their positioning and thus would have low detectability (sensu Schluter and Price 1993; see also Tazzyman et al. 2014). The low detectability likely hinders the effectiveness of pseudo-tail spots as signals for signal receivers (and eavesdroppers), potentially explaining the lack of fixation of this trait.

In summary, some Asian barn swallows exhibit pseudo-tail spots in their undertail coverts, which closely resemble white tails spots. The occurrence and persistence of pseudo-tail spots suggests the potential for exaggeration of existing sexual signals (i.e., white tail spots, here). Unlike theoretical prediction, the spread of pseudo-tail spots appears slow, possibly due to their low detectability. This makes the white spots on both the tail and undertail coverts of barn swallows a rare opportunity to directly observe how costly sexual signal can exaggerate across traits in the wild.

## ACKNOWLEDGMENTS

I am grateful to the residents of Miyazaki Prefecture for their kind support and assistance. I also thank Dr. Yutaka Nakamura, Sunao Suzuki, and Toyofumi Sueyoshi for preliminary field surveys and valuable information regarding swallows there. The manuscript benefited from comments by Drs Emi Arai and Keita Tanaka. I would like to thank Dr. Nobuyuki Kutsukake and the laboratory members of the Graduate University for Advanced Studies. I also thank Dr. Motoomi Yamaguchi for his support. This study is partially funded by Japan Bird Research Association and Kyoko Kajimoto, Naoko Endo, Yusuke Sawa, Hideki Osaka, Naoko Eimura, Tatsuo Sato, Hitoshi Watanabe, Mutsuyuki Ueta, Nanae Kato, Kentaro Takagi, Kazuo Koyama and anonymous supporters. I also thank anonymous referees for their comments on the manuscript.

---

**Fig. S1.**
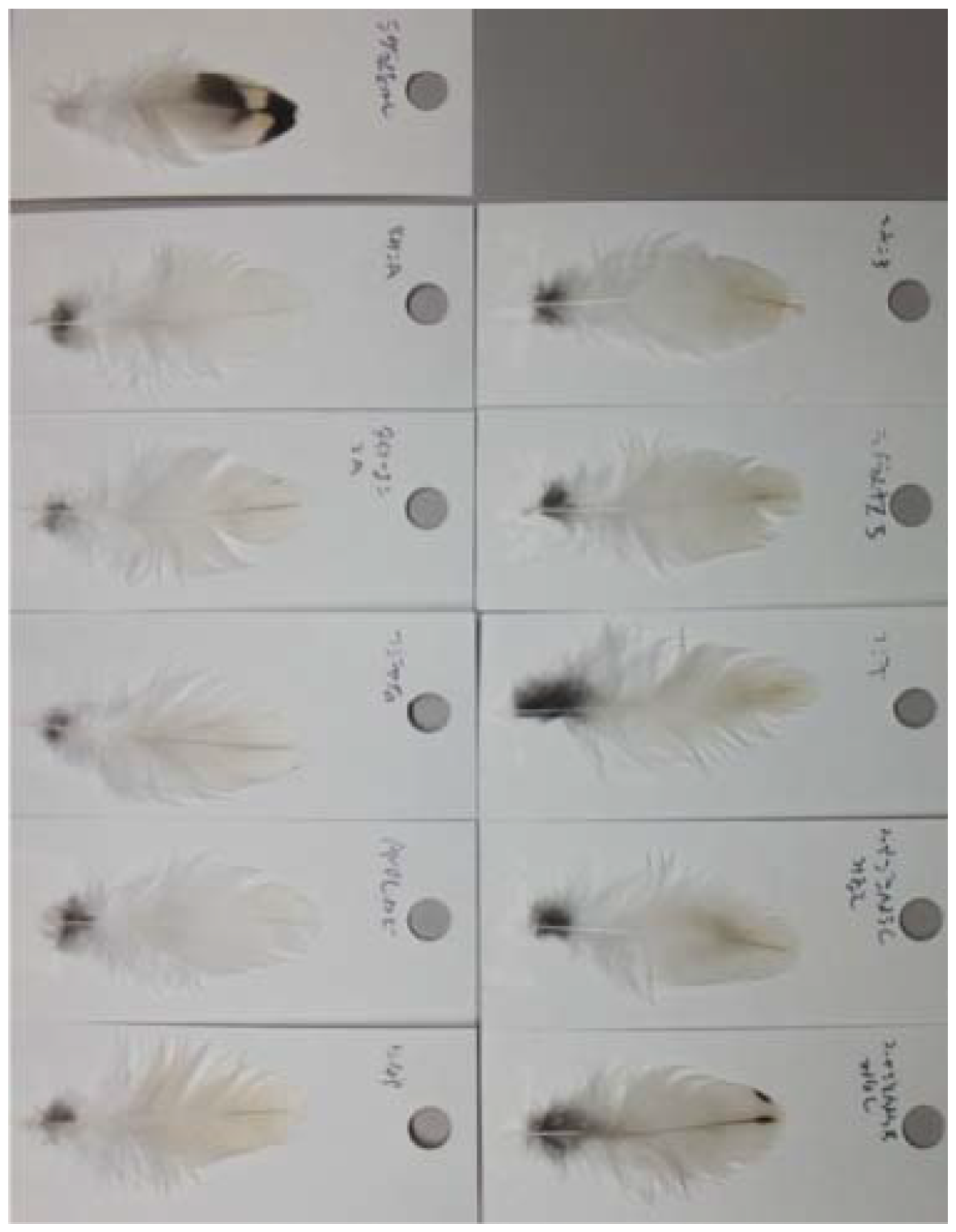
Single undertail coverts feathers plucked from the focal male (top left) and ten additional swallows from Miyazaki Prefecture, 2014. Left: feathers from males. Right: feathers from females.

**Fig. S2.**
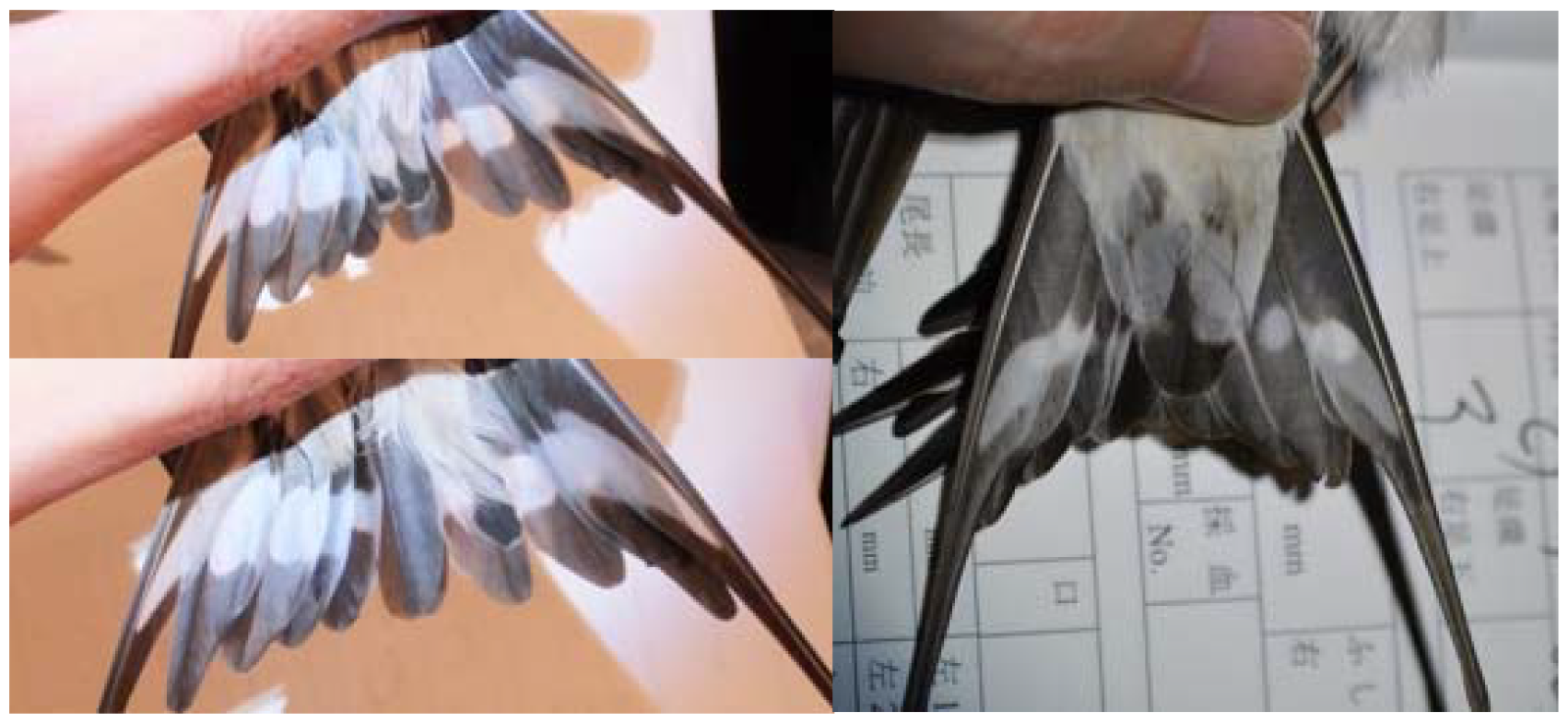
Additional examples of pseudo-tail spots in Asian barn swallows, *Hirundo rustica gutturalis*. Top left: male, Hayama population, 2015. Bottom left: the same individual as upper left. Right: female, Miyazaki population, 2024. Note that central tail feathers lack white spots.

**Fig. S3.**
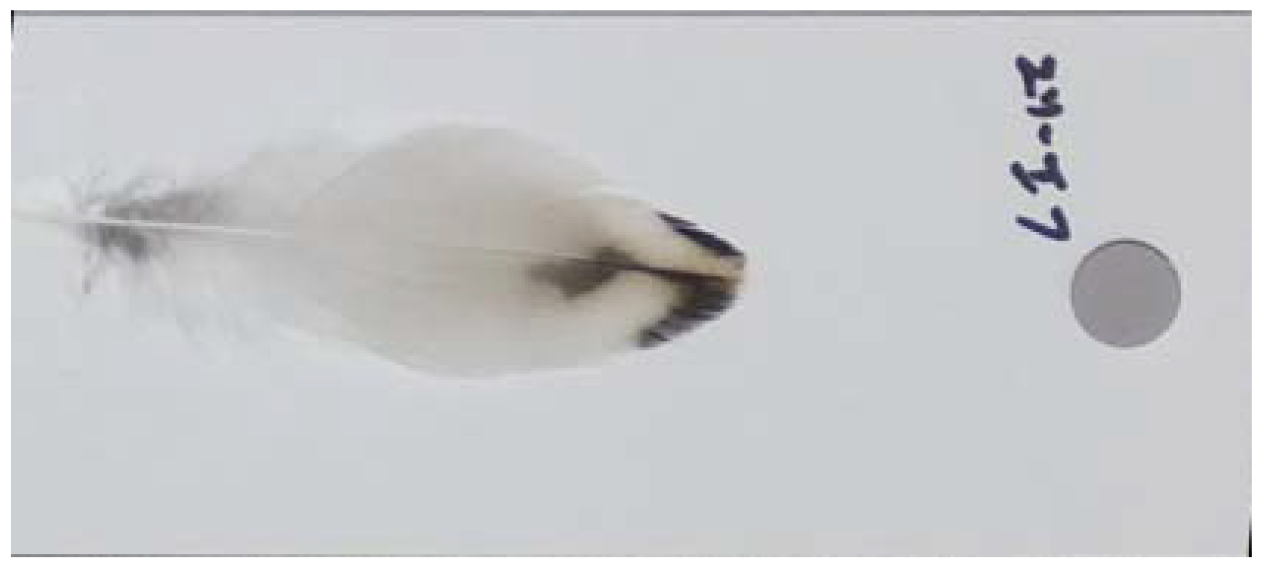
Single undertail covert feather from a female barn swallow in the Hayama population, Kanagawa Prefecture, 2014.

## Notes

### Competing Interest Statement

The authors have declared no competing interest.

